# Diversity of carbapenem-resistant *Acinetobacter baumannii* and bacteriophage-mediated spread of the Oxa23 carbapenemase

**DOI:** 10.1101/2020.09.08.287599

**Authors:** Alaa Abouelfetouh, Jennifer Mattock, Dann Turner, Erica Li, Benjamin A. Evans

## Abstract

Carbapenem-resistant *A. baumannii* are prevalent in low- and middle-income countries such as Egypt, but little is known about the molecular epidemiology and mechanisms of resistance in these settings. Here we characterise carbapenem-resistant *A. baumannii* from Alexandria, Egypt, and place it in a regional context. 54 carbapenem-resistant isolates from Alexandria Main University Hospital, Egypt, collected between 2010 and 2015 were genome sequenced using Illumina technology. Genomes were *de novo* assembled and annotated. Genomes for 36 isolates from the Middle East region were downloaded from GenBank. Core gene compliment was determined using Roary, and analyses of recombination were performed in Gubbins. MLST sequence type and antibiotic resistance genes were identified. The majority of Egyptian isolates belonged to one of 3 major clades, corresponding to Pasteur MLST clonal complex (CC^PAS^) 1, CC^PAS^2 and sequence type (ST^PAS^) 158. Strains belonging to ST^PAS^158 have been reported almost exclusively from North Africa, the Middle East and Pakistan, and may represent a region-specific lineage. All isolates carried an *oxa23* gene, six carried *bla*_NDM-1_, and one carried *bla*_NDM-2_. The *oxa23* gene was located on a variety of different mobile elements, with *Tn2006* predominant in CC^PAS^2 strains, and *Tn2008* predominant in other lineages. Of particular concern, in 8 of the 11 CC^PAS^1 strains, the carbapenemase gene was located in a temperate bacteriophage phiOXA, previously identified only once before in a CC^PAS^1 clone from the US military. The carbapenem-resistant *A. baumannii* population in Alexandria Main University hospital is very diverse, and indicates an endemic circulating population, including a region-specific lineage. The major mechanism for *oxa23* dissemination in CC^PAS^1 isolates appears to be a bacteriophage, presenting new concerns about the ability of these carbapenemases to spread throughout the bacterial population.

**Data Summary:** The whole genome shotgun sequences of the isolates from this study have been deposited at DDBJ/ENA/GenBank under the BioProject accession number PRJNA659545. The individual genome accession numbers for each isolate are as follows: A1a, JACSUC000000000; A2, JACSUB000000000; A4, JACSVQ000000000; A5, JACSUA000000000; A6, JACSTZ000000000; A7-T, JACSVP000000000; A8-T, JACSVO000000000; A8a, JACSTY000000000; A9, JACSTX000000000; A10, JACSTW000000000; A10a, JACSTV000000000; A11a, JACSTU000000000; A13a, JACSTT000000000; A14a, JACSTS000000000; A15, JACSTR000000000; A16, JACSTQ000000000; A18, JACSTP000000000; A21, JACSVN000000000; A22, JACSTO000000000; A27, JACSTN000000000; A30, JACSTM000000000; A31, JACSTL000000000; A34, JACSTK000000000; A35, JACSTJ000000000; A36, JACSTI000000000; A39, JACSTH000000000; A40, JACSTG000000000; A41, JACSTF000000000; A42, JACSTE000000000; A43, JACSTD000000000; A44, JACSTC000000000; A45, JACSTB000000000; A46, JACSTA000000000; A64, JACSSZ000000000; A68, JACSSY000000000; A69, JACSSX000000000; A70, JACSSW000000000; A71, JACSVM000000000; A72, JACSSV000000000; A73, JACSSU000000000; A74, JACSST000000000; A75, JACSSS000000000; A78, JACSSR000000000; A82, JACSSQ000000000; A83, JACSVL000000000; A84, JACSSP000000000; A85, JACSSO000000000; A86, JACSVK000000000; A87, JACSSN000000000; A88, JACSSM000000000; A89, JACSSL000000000; A92, JACSSK000000000; A5910, JACSSJ000000000; A6135, JACSVJ000000000.

## Introduction

The bacterium *Acinetobacter baumannii* is a major opportunistic hospital-acquired pathogen that is listed by the World Health Organization (WHO) as in critical need of new treatment options due to its multidrug-resistant nature (1). In particular, the frequency of carbapenem-resistant *A. baumannii* has been steadily increasing over the last two decades, leaving very few treatment options available to combat this pathogen (2). However, carbapenem-resistant *A. baumannii* are not uniformly distributed across the globe, with higher rates of resistance found in low- and middle-income countries (3–6), though rates in some southern and eastern European countries have now also reached very high levels (7). In countries in the Middle East and North Africa high levels of carbapenem-resistant *A. baumannii* are reported, with frequencies of 70% of isolates or greater being common (8). Despite these very high rates of resistance, there are relatively few studies investigating the molecular epidemiology of the antibiotic-resistant strains.

Carbapenem resistance in *A. baumannii* is usually the result of the expression of an OXA-type β-lactamase, or occasionally metallo-β-lactamases such as the IMP, VIM and NDM groups (9). The acquired OXA-type β-lactamases in *A. baumannii* are encoded by genes belonging to five main groups – *oxa23* (or *bla*_OXA-23-like_), *oxa40* (or *bla*_OXA-40-like_), *oxa58* (or *bla*_OXA-58-like_), *oxa134* (or *bla*_OXA-134-like_), and *oxa143* (or *bla*_OXA-143-like_) (10, 11). In addition, all *A. baumannii* carry an intrinsic OXA β-lactamase gene called *oxaAb* (or *bla*_OXA-51-like_), certain alleles of which when highly expressed due to the presence of an *ISAba1* insertion sequence upstream can confer carbapenem resistance (12–14). The most common of these resistance mechanisms globally is *oxa23* (15). In Egypt, and other countries in the region, *oxa23* is so prevalent it can be found in up to 100% of carbapenem-resistant isolates, with frequencies greater than 70% being the norm (16–19). In *A. baumannii oxa23* is usually located on a transposon mobilised by one or more insertion sequences (IS), which has enabled the resistance gene to be spread to many different plasmids and many different lineages within the species (20). Despite the particularly high prevalence of *oxa*23 in low- and middle-income countries such as Egypt, there are very few studies that have investigated the mobile genetic elements carrying the gene which is crucial to gaining an understanding of the local population genetics of the species.

The majority of *A. baumannii* isolates belong to one of eight International Clones (ICs), which correspond to specific multi-locus sequence typing (MLST) sequence types (STs) and clonal complexes (CCs) (15, 21). There are two MLST schemes for *A. baumannii* – the Pasteur scheme (22) and the Oxford scheme (23) with the Pasteur scheme containing genes that are less prone to recombination than the those in the Oxford scheme (24). Globally, isolates belonging to IC2, corresponding to CC^PAS^2, are the most common, though there are exceptions such as Latin American countries where isolates belonging to IC4 (CC^PAS^15), IC5 (CC^PAS^79) and IC7 (CC^PAS^25) are predominant (25, 26). In many low- and middle-income countries MLST is too costly to perform on large numbers of isolates, and so at present we often rely on a small number of studies to provide an indication of what the national epidemiology may be. In Egypt, studies have indicated that CC^PAS^2 is the most common CC, but that a large number of isolates from other CCs or that don’t belong to any of the defined CCs make up a substantial portion of the population (16, 18, 19, 27). The aim of our study was to define the local population structure of *A. baumannii* in Alexandria Main University Hospital, Egypt, and identify the mobile genetic elements responsible for resistance gene dissemination.

## Methods

### Bacterial isolates and antimicrobial susceptibility testing

A total of 54 carbapenem resistant *A. baumannii* clinical isolates obtained from patients presenting at Alexandria Main University Hospital (AMUH) between 2010 and 2015 were included in the study. This is the largest hospital in the northern sector of Alexandria and a major referral hospital. The isolates were identified by conventional methods, including colony morphology, aerobic growth at 44°C on MacConkey agar, and species designations obtained using the Vitek system (bioMérieux, UK). The identity of the isolates was further confirmed by PCR amplification of the intrinsic *oxaAb* (*bla*_OXA-51-like_) gene as well as Matrix Assisted Laser Desorption/Ionization - Time of Flight Mass Spectrometry (MALDI-TOF MS) (Bruker Daltonik, USA). The identified isolates were stored at −80°C prior to subsequent characterisation (28). The susceptibility of the isolates to 5 antibiotics was determined using agar dilution or broth microdilution (for colistin) and the results were interpreted according to the Clinical and Laboratory Standards Institute guidelines (2018)(29). The antibiotics used were imipenem, meropenem, colistin, levofloxacin and amikacin.

### Whole genome sequencing and analyses

Genomic DNA was extracted using the Wizard® Genomic DNA Purification Kit (Promega) according to the manufacturer’s instructions. A Qubit fluorometer (Life Technologies) was used to quantify the extracted DNA. Dual indexing library preparation was carried out using the Nextera XT DNA Preparation Kit (Illumina Inc., San Diego, CA, USA). Whole genome sequencing of the library was performed on an Illumina MiSeq using the 2 x 250 bp paired-end protocol. Following quality filtering of the reads using Trimmomatic v0.36 (30) and FastQC v0.11.5 (31), genomes were *de novo* assembled with Spades v3.11.1 (32), and annotated using Prokka v1.11 (33). The assemblies were quality checked using QUAST (34). The Sequence Read Archive was searched using keywords of Middle Eastern countries, and the genomes of an additional 36 strains were downloaded and included in subsequent analyses. The genomes of a further 17 geographically and genomically diverse strains (35) were also downloaded and included in subsequent analyses. The core genome content of the strain collection was determined using Roary v3.12.0 (36), and the core gene phylogeny estimated using FastTree v2.1.10 (37). Isolates A8-T and A74 were chosen to be the reference genomes for CC^PAS^1 and CC^PAS^2 respectively for subsequent variant calling. Sequences belonging to CC^PAS^1 and CC^PAS^2 were mapped to the reference genomes and variant called using PHEnix v1.3 (38). A SnapperDB v1.0.6 for each CC was created, allowing inclusion of SNPs with a minimum average read depth of 10 (39). Whole genome alignments were generated including isolates 10,000 SNPs from the CC1 reference and 20,000 SNPs from the CC2 reference, which were used as input for Gubbins. Estimates of recombination within clades identified in the phylogeny were conducted with Gubbins v2.3.1 using default settings (40). The Pasteur MLST sequence type (ST) of each isolate was determined from the whole genome sequence using the online Center for Genomic Epidemiology’s MLST software (41), and antibiotic resistance genes using ARIBA (42) using the CARD (43) and SRST2 (44) databases. All *oxaAb* alleles were confirmed using the BLAST function on the Beta-lactamase database (11)(http://www.bldb.eu/). Analyses of the accessory genome were conducted using PANINI (45).

### Annotation of phiOXA-A35

The phiOXA-A35 sequence was constructed by alignment of three contigs and manual resolution of overlaps from sequence/assembly data obtained for *A. baumannii* A35. phiOXA open reading frames were initially annotated using PROKKA v1.12 and then refined using BLASTp, InterProScan (46) and HHpred (47). Prediction of tRNAs was performed using tRNAscan-SE 2.0 (48). Alignments of the portal vertex and major capsid proteins were performed using Clustal Omega (49) and phylogenetic trees constructed using IQTree v1.6.12 with ModelFinder, SH-aLRT test and ultrafast bootstrap with 1,000 replicates (50–52). Read coverage of phiOXA-A35 was calculated using QualiMap v.2.2.2 (53).

### *Genetic environment of* oxa23

In the program Geneious R10 (https://www.geneious.com/), for each *oxa23*-positive strain the contig containing *oxa23* was identified, then all these contigs were aligned. The alignment was used to group strains by similarity of the sequence surrounding *oxa23*. Where appropriate, the presence of insertion sequences *ISAba1* and *ISAba125* surrounding the *oxa23* gene were confirmed by PCR using combinations of primers ISAba1-B (54), OXA-23-F and OXA-23-R (55), and ISAba125-F (5’-TAAAACTATTCATGAGCGCC-3’). To obtain the complete sequences of the prophages containing *oxa23*, contigs were aligned against the phiOXA sequence from strain AB5075-UW (accession no. CP008706.1). PCRs with primers Phi-F (5’ – CGT TGT TGG GCT TCT AGT GC – 3’) and OXA-23-R (55) were used to confirm the contig joins either side of the IS*Aba1* insertion sequence.

### Bacteriophage induction

Bacterial cultures were grown overnight in LB at 37°C and shaking at 180 rpm. Overnight cultures were diluted to an OD_600_ of approximately 0.05 using pre-warmed LB, then incubated at 37°C and 180 rpm until the OD_600_ reached 0.2. Cultures were then divided to generate two treatment cultures and two control cultures per strain. Mitomycin C was added to a final concentration of 2 μg/mL to the treatment cultures, then culture tubes containing mitomycin C were wrapped in foil to block out light, then treatment and control cultures were incubated at 37°C. The OD_600_ of cultures was recorded every 30 minutes to identify a marked drop in the optical density in the mitomycin C-treated cultures, representing induction. Once this was observed, all cultures were centrifuged at 10,000 g for 5 minutes, filter sterilised through a 0.22 μm filter, and treated with DNase (TURBO DNase, Invitrogen) and RNase A (Thermo Scientific) to remove all bacteria and free nucleic acid from the cell lysate. The presence of intact bacteriophage carrying *oxa23* in the bacterial cell lysate was determined by PCR using primers OXAphi-F (5’ – GGAAATGCGGTCAGAAATGC – 3’) situated within *oxa23* and OXAphi-R (5’ – TGGACCCTGTAGATTTTGCC – 3’) situated within a phage tail protein gene, giving a 1,032 bp product size. PCR conditions were 95°C for 10 minutes, followed by 40 cycles of 95°C for 30 seconds, 55°C for 30 seconds, and 72°C for 1 minutes, with a final extension of 72°C for 5 minutes. A 1 μL culture volume from a phiOXA-positive strain was used as a positive control.

## Results

Analysis of the antibiotic susceptibilities of the Egyptian isolates showed that, as expected due to the isolates being selected for their carbapenem resistance, they were all resistant to imipenem and meropenem (Table 1). Furthermore, all isolates were found to be multidrug resistant as they were all resistant to both levofloxacin and amikacin (Table 1)(29). There were also high levels of resistance to colistin, with the MIC for 24 isolates (44%) above the breakpoint of 2 mg/L. The majority of these were only one dilution above the breakpoint i.e. 4 mg/L. However, 7 isolates (13%) had a very high colistin MIC of 256 mg/L.

**Table 1:** Antibiotic susceptibility data, MLST assignments, carbapenem resistance genes and associated mobile genetic elements of Egyptian isolates.

| Strain | MICs (mg/L) |  |  |  |  | MLST (Pasteur) |  | Antibiotic resistance genes |  | *oxa23* mobile element<sup>1</sup> |
| --- | --- | --- | --- | --- | --- | --- | --- | --- | --- | --- |
|  | IMI | COL | MER | LEV | AMI | ST | CC | *oxaAb* |  |  |
| 1a | 4 | 4 | 32 | 256 | >512 | 664 | 2 | 66 | *oxa23* | A |
| 2 | 8 | 4 | 64 | 8 | 64 | 1 | 1 | 69 | *oxa23, bla<sub>NDM-1</sub>* | nd |
| 4 | 16 | 4 | 256 | 16 | 128 | 1 | 1 | 69 | *oxa23* | nd |
| 5 | 32 | 4 | 64 | 32 | >512 | 25 | 25 | 64 | *oxa23* | A |
| 6 | 8 | 2 | 32 | 16 | >512 | 1 | 1 | 69 | *oxa23* | phiOXA |
| 7-T | 8 | 1 | 64 | 16 | >512 | 1 | 1 | 69 | *oxa23* | phiOXA |
| 8-T | 8 | 1 | 64 | 16 | 128 | 1 | 1 | 69 | *oxa23* | phiOXA |
| 8a | 16 | 2 | 32 | 32 | >512 | 15 | 15 | 51 | *oxa23* | A |
| 9 | 8 | 2 | 64 | 16 | 128 | 158 | 158 | 65 | *oxa23* | H |
| 10 | 8 | 2 | 32 | 16 | 128 | 158 | 158 | 65<sup>2</sup> | *oxa23* | C |
| 10a | 16 | 4 | 32 | 32 | >512 | 2 | 2 | 66 | *oxa23* | A |
| 11a | 16 | 2 | 32 | 256 | 64 | 1535 |  | 65<sup>3</sup> | *oxa23* | E |
| 13a | 4 | 2 | 32 | 256 | >512 | 664 | 2 | 66 | *oxa23* | A |
| 14a | 16 | 2 | 16 | 256 | >512 | 664 | 2 | 66 | *oxa23* | A |
| 15 | 8 | 4 | 64 | 16 | >512 | 25 | 25 | 64 | *oxa23* | A |
| 16 | 8 | 4 | 64 | 8 | 64 | 85 | - | 94 | *oxa23* | nd |
| 18 | 8 | 2 | 64 | 32 | >512 | 19 | 1 | 69 | *oxa23* | J |
| 21 | 32 | 4 | 64 | 32 | 128 | 1 | 1 | 69 | *oxa23, bla<sub>NDM-1</sub>* | phiOXA |
| 22 | 8 | 4 | 32 | 32 | >512 | 2 | 2 | 66 | *oxa23* | A |
| 27 | 8 | 4 | 32 | 256 | 512 | 2 | 2 | 66 | *oxa23* | F |
| 30 | 8 | 4 | 32 | 256 | >512 | 158 | 158 | 65 | *oxa23* | nd |
| 31 | 16 | 2 | 64 | 256 | >512 | 15 | 15 | 51 | *oxa23* | G |
| 34 | 8 | 4 | 64 | 128 | >512 | 158 | 158 | 65 | *oxa23* | nd |
| 35 | 4 | 4 | 64 | 128 | 64 | 1 | 1 | 69 | *oxa23* | phiOXA |
| 36 | 8 | 2 | 64 | 128 | 128 | 158 | 158 | 65 | *oxa23* | nd |
| 39 | 4 | <0.5 | 64 | 16 | 64 | 1 | 1 | 69 | *oxa23* | phiOXA |
| 40 | 32 | 2 | 64 | 128 | >512 | 1 | 1 | 69 | *oxa23, bla<sub>NDM-1</sub>* | nd |
| 41 | 16 | 2 | 32 | 32 | 64 | 1 | 1 | 69 | *oxa23* | phiOXA |
| 42 | 32 | 2 | 64 | 16 | 64 | 158 | 158 | 65 | *oxa23* | B |
| 43 | 16 | 4 | 64 | 16 | 128 | 158 | 158 | 65 | *oxa23* | B |
| 44 | 8 | 4 | 64 | 128 | 512 | 158 | 158 | 65 | *oxa23* | B |
| 45 | 32 | 2 | 32 | 16 | 128 | 158 | 158 | 65 | *oxa23* | nd |
| 46 | 8 | 2 | 16 | 32 | 64 | 1 | 1 | 69<sup>4</sup> | *oxa23* | phiOXA |
| 64 | 8 | 2 | 64 | 256 | >512 | 664 | 2 | 66 | *oxa23* | A |
| 68 | 8 | 2 | 32 | 128 | >512 | 158 | 158 | 65 | *oxa23* | D |
| 69 | 8 | 2 | 32 | 256 | >512 | 2 | 2 | 66 | *oxa23* | C |
| 70 | 16 | 2 | 32 | 128 | 512 | 103 | - | 70 | *oxa23, bla<sub>NDM-2</sub>* | nd |
| 71 | 8 | 2 | 32 | 32 | 512 | 2 | 2 | 66 | *oxa23* | F |
| 72 | 16 | 4 | 64 | 64 | 64 | 2 | 2 | 66 | *oxa23* | A |
| 73 | 8 | 2 | 64 | 64 | >512 | 2 | 2 | 66 | *oxa23* | A |
| 74 | 8 | 2 | 32 | 256 | >512 | 664 | 2 | 66 | *oxa23* | A |
| 75 | 8 | 2 | 16 | 128 | >512 | 2 | 2 | 66 | *oxa23* | F |
| 78 | 8 | 256 | 16 | 128 | 512 | 570 | 2 | + | *oxa23, bla<sub>NDM-1</sub>* | A |
| 82 | 64 | 256 | 64 | 128 | >512 | 2 | 2 | 66 | *oxa23* | C |
| 83 | 64 | 256 | 64 | 16 | >512 | 2 | 2 | 66 | *oxa23* | A |
| 84 | 64 | 2 | 64 | 16 | >512 | 2 | 2 | 66 | *oxa23* | A |
| 85 | 64 | 256 | 32 | 16 | >512 | 2 | 2 | 66 | *oxa23* | A |
| 86 | 64 | 256 | >256 | 128 | >512 | 570 | 2 | 66 | *oxa23, bla<sub>NDM-1</sub>* | A |
| 87 | 64 | 256 | 64 | 16 | 16 | 1 | 1 | 69 | *oxa23* | I |
| 88 | 64 |  | 64 | 128 | >512 | 2 | 2 | 66 | *oxa23* | A |
| 89 | 64 | 256 | 64 | 128 | >512 | 2 | 2 | 66 | *oxa23* | F |
| 92 | 32 | 4 | 64 | 128 | >512 | 2 | 2 | 66<sup>5</sup> | *oxa23* | A |
| 5910 | 128 | 2 | >256 | 128 | >512 | 2 | 2 | 66 | *oxa23* | A |
| 6135 | 8 | 2 | 32 | 32 | >512 | 600 | 2 | 66 | *oxa23, bla<sub>NDM-1</sub>* | A |

<sup>1</sup>nd, not determined – the Illumina sequence data was not able to resolve contigs showing the genetic environment of *oxa23* in these isolates. <sup>2</sup>Contig break giving incomplete gene, with 243/243 amino match to *oxaAb*(65). <sup>3</sup>Contig break giving incomplete gene, with 265/266 amino acid match to *oxaAb*(65). <sup>4</sup>Contig break giving incomplete gene, with 266/266 amino acid match to *oxaAb*(69). <sup>5</sup>Contig break giving incomplete gene, with 266/266 amino acid match to *oxaAb*(66). +, *oxaAb* gene was not identified in genome sequence but was positive by PCR.

The majority of Egyptian isolates belonged to one of three major well-supported clades based upon core gene sequences, corresponding to Pasteur MLST clonal complex (CC^PAS^) 1 (13 isolates), CC^PAS^2 (24 isolates) and ST^PAS^158 (10 isolates) (Table 1, Fig. 1). CC^PAS^1 isolates belong to International Clone (IC) 1, while CC^PAS^2 isolates belong to IC2 (21, 56). In addition, two isolates belonged to ST^PAS^15, which are members of IC4, and two isolates belonged to ST^PAS^25, which are members of IC7 (Table 1). All isolates belonging to the ICs carried the *oxaAb* allele previously shown to be associated with their respective IC, with isolates in CC^PAS^1 (IC1) carrying *oxaAb*(69), isolates in CC^PAS^2 (IC2) carrying *oxaAb*(66), isolates in ST^PAS^15 (IC4) carrying *oxaAb*(51), and isolates in ST^PAS^25 (IC7) carrying the *oxaAb*(64) allele (57). Isolates in ST^PAS^158 carried *oxaAb*(65) alleles, which are usually associated with CC^PAS^79 and IC5. However, it should be noted that the *oxaAb*(65) allele in the ST^PAS^158 isolates differed from the original *oxaAb*(65) allele (GenBank accession no. AY750908) by 3 silent substitutions (T90C, C636T and A663G). We compared our ST^PAS^158 isolates with other published or publicly available data, which demonstrated that this particular *oxaAb*(65) variant is a feature of ST^PAS^158 isolates in general (Table 2), and is distinct from CC^PAS^79 (IC5) isolates. Isolates from CC^PAS^1 and CC^PAS^2 were more diverse than ST^PAS^158 isolates. This was evident in gene conservation analysis with 3,069 genes shared by 90% of ST^PAS^158 isolates, whereas only 2,394 genes were shared by 90% of CC^PAS^2 isolates, and 2,600 genes shared by 90% of CC^PAS^1 isolates. For all 3 of the major clades identified, the isolates from AMUH did not form their own specific sub-clades, but were interspersed with the strains both from other Middle Eastern countries as well as with the globally distributed strains (Figure 1). Interestingly a similar pattern was observed with respect to the accessory genome (Figure 2). Based upon their accessory genomes, CC^PAS^1, CC^PAS^2 and ST^PAS^158 isolates clustered together. The only exception was isolate 11a which clustered with the CC^PAS^2 isolates (the grey dot found on the right-hand edge of the orange CC^PAS^2 cluster in Figure 2A). However, this is not too surprising given that of all the non-CC^PAS^2 isolates, 11a is most closely related to CC^PAS^2 at the core genome level (Figure 1). The accessory genome clusters did not show any geographic signal (Figure 2B), in agreement with the lack of geographical signal in the core gene tree (Figure 1). Together these data demonstrate two points: firstly that there are both multiple circulating clonal lineages, and multiple circulating sub-lineages within each clonal lineage, that are responsible for infecting patients in AMUH, and secondly that while the accessory genome is shared across isolates from several different countries within a clonal lineage, there is little sharing of the accessory genome between clonal lineages.

**Figure 1:**
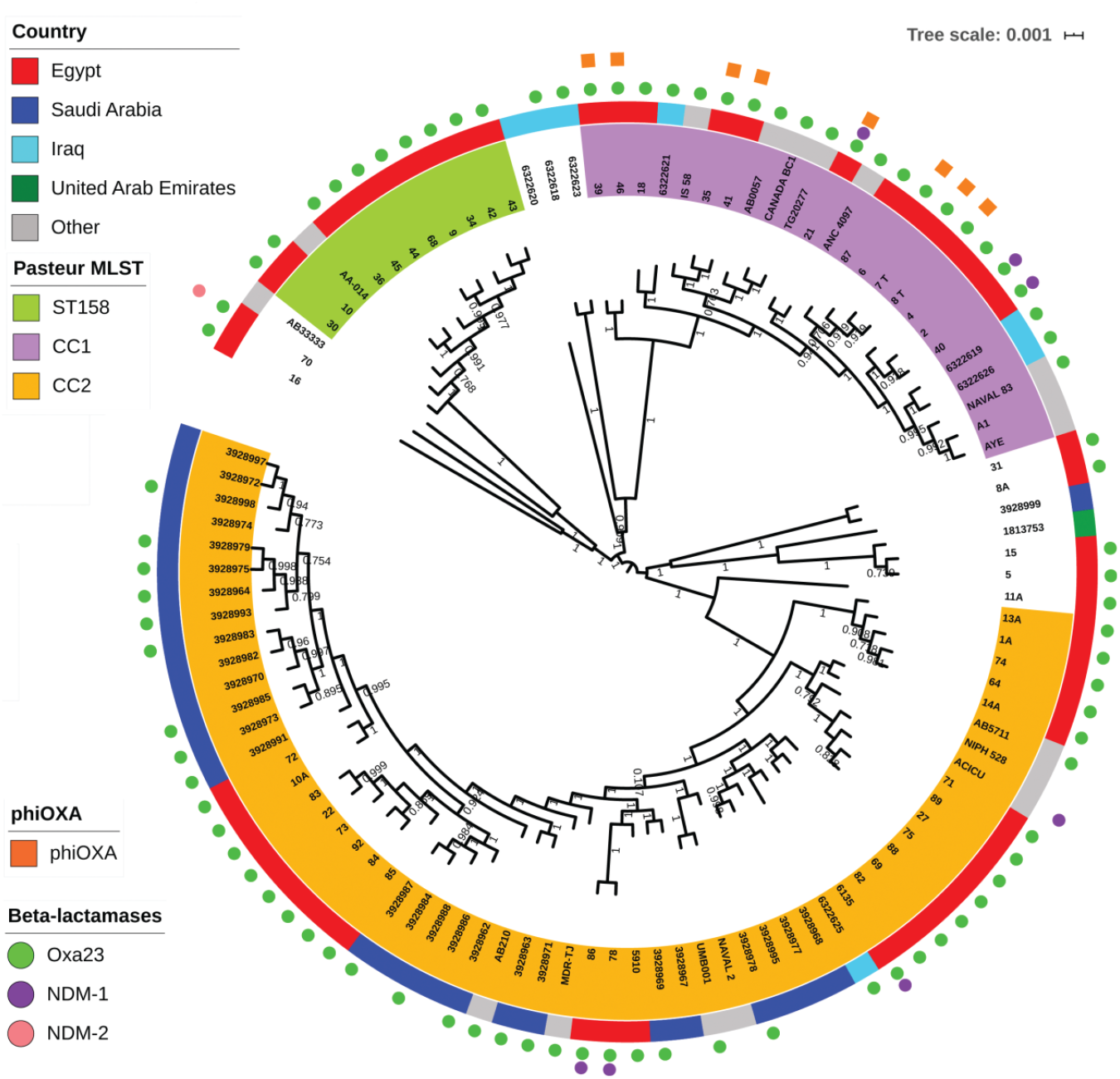
Core gene tree of all isolates. In the centre is the core gene tree generated in FastTree (37) using a core gene alignment output from Roary (36). The tree is scaled by genetic distance, and branch labels indicate level of support based upon the Shimodaira-Hasegawa test using 1,000 resamples. Leaves are labelled with isolate names or SRA accession numbers, and are colour coded to highlight the three major Pasteur MLST scheme clonal complex or sequence types identified in this study. The ST/CC of isolates that are not coloured can be seen in Table 1. The outer solid coloured ring indicates the geographic source of the isolates. The outer rings of shapes indicate β-lactamases and phiOXA encoded by the isolates.

**Figure 2:**
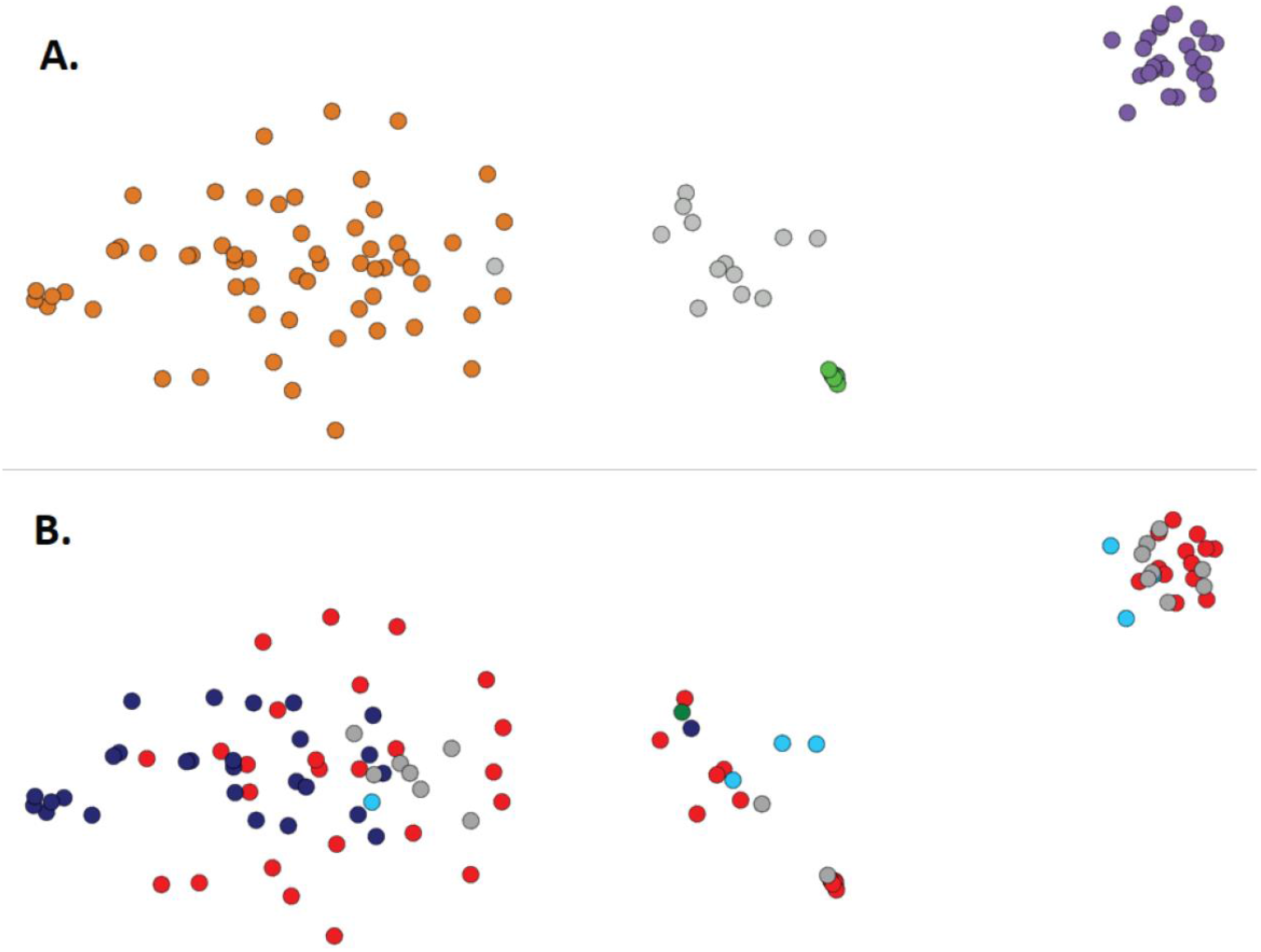
Clustering of isolates by similarity of their non-core genomes using PANINI. Each dot represents an isolate, and the distance between isolates indicates the similarity of their accessory genomes. (A) the network is coloured according to the MLST data as in figure 1 (CC1 is purple, CC2 is orange, ST158 is green, and other STs are grey). (B) the network is coloured according to the country of origin of the isolates as in Figure 1 (Egypt is red, Saudi Arabia is dark blue, Iraq is light blue, the UAE is green, and other countries outside the Middle East are grey).

**Table 2:** CC^PAS^158 and CC^OX^499 isolates reported in the literature or in public databases.

| Isolate<sup>a</sup> | ST<sup>PAS</sup> | ST<sup>OX</sup> | Country | Year | *oxaAb*<sup>b</sup> | Accession no. | Ref. |
| --- | --- | --- | --- | --- | --- | --- | --- |
| 10 isolates | 158 | 499<sup>f</sup> | Egypt | 2010-15 | 65* |  | This study |
| 1309; 2226C | 158 | - | Turkey | 2009 | - | - | PubMLST |
| 2313; AA-014 | 158 | 960 | Iraq | 2008 | 65* | GCA_000335595 | (86) |
| 3826; 778944; ABC002 | 158 | 1717 | Egypt | 2012 | - | - | PubMLST |
| K50 | 158 | - | Kuwait | 2008 | 65* | OHJL000000000 | (87) |
| Unnamed | 158 | - | Lebanon | 2013 | 65* | - | (88) |
| Ab-Pak-Pesh-01 | 158 | - | Pakistan | 2015 | 65* | SMUB010000000 | (64) |
| Ab-Pak-Pesh-07 | 158 | - | Pakistan | 2015 | 65* | QQPV000000000 | (64) |
| Ab-Pak-Pesh-28 | 158 | - | Pakistan | 2015 | 65* | QQPZ000000000 | (64) |
| AMA 341 | 158 | 499 | Denmark<sup>c</sup> | 2012 | 65<sup>d</sup> | SAMN03160609 | (89) |
| Two isolates | 158 | - | Kuwait | 2011-12 | - | - | (90) |
| ACB69C | 158 | - | Turkey | 2009-11 | - | - | (91) |
| 30 isolates | 158 | - | Kuwait | 2007-08 | 66 | - | (92) |
| 7 isolates | 158 | 499 | Tunisia | 2008-09 | - | - | (93) |
| 1830; J17 | 342 | - | China | 2011 | - | - | (94) |
| 3840; ACIN00151 | 342 | 1776 | USA | 2016 | 694 | PubMLST<sup>e</sup> | PubMLST |
| 2178; A.baumannii64 | 615 | - | Egypt | 2012 | - | - | (16) |
| 2180; A.baumannii85 | 615 | - | Egypt | 2013 | - | - | (16) |
| 2182; A.baumannii108 | 618 | - | Egypt | 2013 | - | - | (16) |
| 3950; TR112 | 1241 | - | Turkey | 2016 | - | - | PubMLST |
| 8 isolates | - | 499 | Egypt | 2015 | - | - | (27) |
| 2 isolates | - | 499 | Saudi Arabia | 2011-13 | - | - | (19) |
| 1 isolate | - | 499 | Kuwait | 2011-13 | - | - | (19) |

<sup>a</sup>If an isolate is known by more than one name, all are provided separated by semicolons. <sup>b</sup>The *oxaAb*(65\*) alleles differ from the original *oxaAb*(65) sequence by 3 silent substitutions. <sup>c</sup>This isolate was likely imported from Egypt. <sup>d</sup>Authors do not state whether the nucleotide sequence differs from the original *oxaAb*(65) sequence. <sup>e</sup>This genome is available through the PubMLST website. <sup>f</sup>One isolate did not have its ST<sup>OX</sup> determined.

All 54 isolates carried an *oxa23* gene (Table 1). In addition, 6 isolates also carried a *bla*_NDM-1_ gene, and one isolate carried *bla*_NDM-2_. The *bla*_NDM_ genes were not clustered in one particular bacterial sequence type, with 3 of the *bla*_NDM-1_ genes located in CC^PAS^1 isolates (in one ST) while the other 3 were located in CC^PAS^2 isolates (across 2 STs) (Table 1, Figure 1). The *oxa23* gene was located on a variety of different mobile genetic elements, with 11 different structures identified (Table 1, Figure 3). Several of these structures were found in multiple isolates: structure A, representing *Tn2006* (58), was the most common and was found in 21 isolates, 18 of which belonged to CC^PAS^2, 2 belonged to CC^PAS^25, and one to CC^PAS^15; structure B was found exclusively in three ST^PAS^158 isolates; structure C in one ST^PAS^158 and two CC^PAS^2 isolates; and structure F in four CC^PAS^2 isolates and appeared to be borne on the chromosome (Table 1, Figure 3). Of particular concern, in 8 of the 13 CC^PAS^1 strains carrying *oxa23*, the carbapenemase gene was located in prophage called phiOXA. This prophage has been identified only once before in the CC^PAS^1 isolate AB5075-UW from the US military (59). In order to determine whether phiOXA can form viable viral particles that contain the *oxaAb* gene, four isolates encoding phiOXA (A8-T, A21, A35 and A39) and one isolate that did not (A18) were treated with mitomycin C to induce bacteriophage, followed by DNase and RNase treatment to remove any DNA that is not contained within a virus particle. Then a PCR for *oxa23*, with an extended initial denaturation phase to lyse bacteriophage particles, was used to identify the carriage of the antibiotic resistance gene by the bacteriophage. Cultures of three of the four isolates tested (A8-T, A35 and A39) that had been treated with mitomycin C were found to have produced intact bacteriophage carrying *oxa23*. No PCR products for *oxa23* were detected for these strains when they were not induced, nor for isolate A18 (phiOXA negative) with either the presence or absence of mitomycin C treatment. These data demonstrate that the phiOXA prophage in these isolates can be induced and form intact bacteriophage particles, and that these bacteriophages carry the *oxa23* gene.

**Figure 3:**
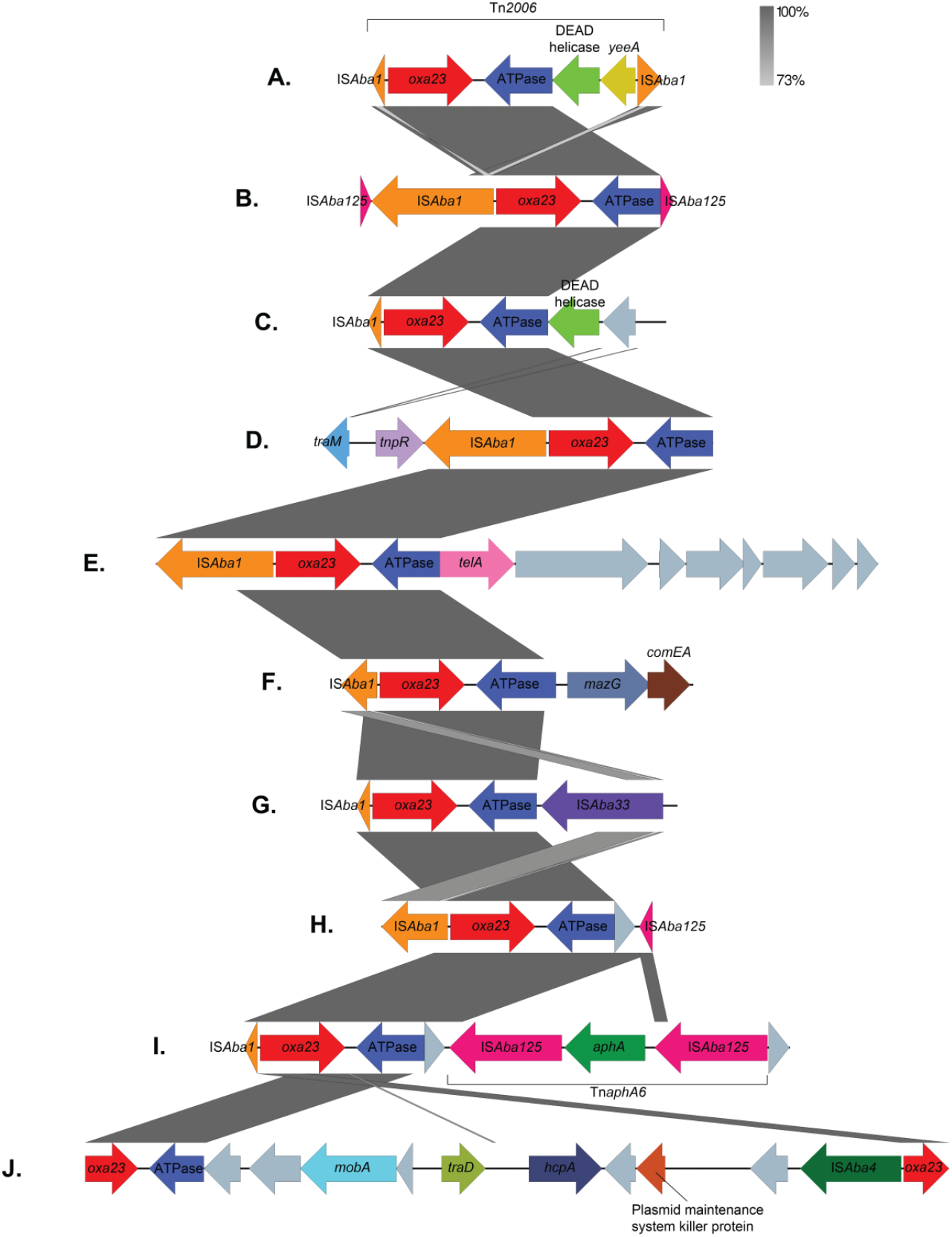
Genetic environments surrounding *oxa23* genes. Arrows represent genes, which are colour-coded by their type. Unlabelled grey genes represent hypothetical proteins. The size of the genes and the distances between them are drawn to scale. Vertical grey boxes indicate homology between sequences ranging between 73% and 100% identity (BLASTn). The diagram was created using Easyfig (95) and annotated in Adobe Photoshop.

Analysis of the sequence of phiOXA-A35 showed it is identical to the bacteriophage/prophage reported in strain AB5075-UW (Figure 4), with mean read coverage of 22 (s.d. = 7). The phiOXA-A35 prophage consists of a contiguous 32 kb region comprising 48 open reading frames (ORFs) with the attL site residing within a tRNA-Leu. The genomic architecture of phiOXA-A35 is similar to that of members of the *Peduovirinae*, a widespread subfamily of temperate bacteriophages that infect γ- and β-proteobacteria and includes *Escherichia* phage P2 and *Pseudomonas* phage phiCTX. The genome can be divided into four modules, representing genes involved in virion morphogenesis and assembly which contains the diagnostic *Q-P-O-N-M-L* capsid gene cluster (60), lysis, replication, and control of lysogeny. This relationship is further supported by phylogenetic analysis of the portal vertex and major capsid protein (Figures S1 and S2). Apart from a single syntenic break representing the three gene cassette containing *ISAba1, oxa23* and a DUF815 domain protein, phiOXA is nearly identical to predicted prophage regions found in *A. baumannii* strains A85, AYE, DA33382, USA15 and WCHAB005078. Comparison of these regions using VIRIDIC (61) suggests that they represent a single species of temperate bacteriophage as each exhibit >95% sequence similarity (62). We propose that phiOXA-A35 represents a new genus within the subfamily *Peduovirinae*. A total of 8 phiOXA ORFs are annotated as hypothetical proteins and whether these represent additional proteins which influence the pathobiology or environmental fitness of their host lysogen remains to be elucidated.

**Figure 4:**
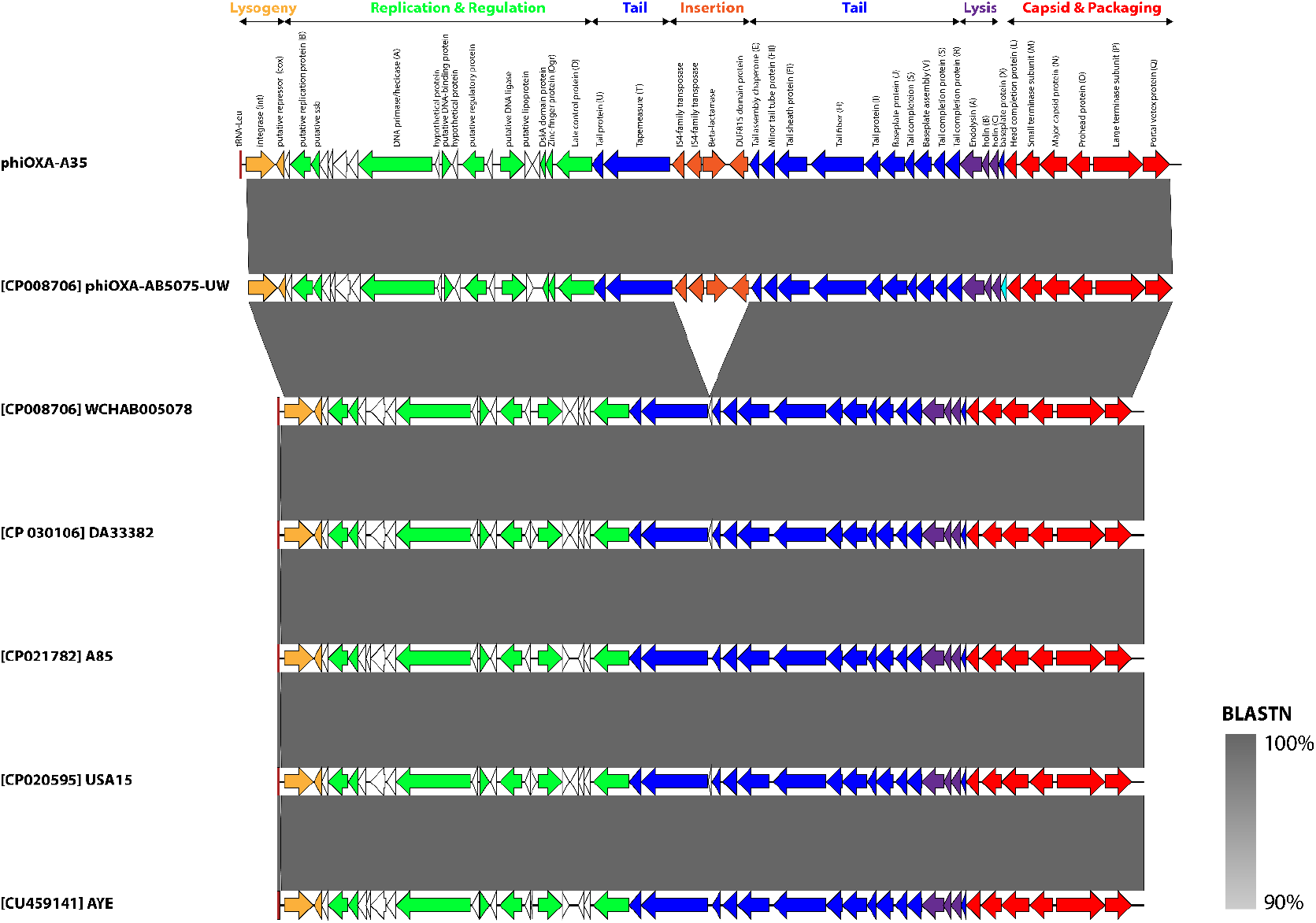
Schematic genome map of phiOXA and related prophages. Prophages are orientated as they appear in their host genome. Arrows depict open reading frames and are coloured according to function. Homologs to gene products in *Escherichia* phage P2 are indicated in parentheses. ORFs encoding hypothetical proteins are shown as black outlines. The tRNA-Leu, representing the attL site, is shown as a dark red rectangle. Shading between entries represents the percent identity (BLASTn) from 90% (light grey) to 100% (dark grey). The map was constructed using Easyfig (95) and annotated in Adobe Illustrator.

## Discussion

In this study we aimed to use genomics to characterise the molecular epidemiology and antibiotic resistance of *A. baumannii* isolates from Alexandria, Egypt. Genome-level studies of this nature from low- and middle-income countries are not common despite the fact that these countries bear the highest burden of antibiotic resistance. By using genomics we can simultaneously characterise antibiotic resistance genes and the genetic environment supporting them, and the fine-scale epidemiological relationships between isolates. It also has the added benefit of being backward-compatible with previous typing methods such as MLST. In the context of Egypt, there are a few studies that have used one of the MLST schemes for *A. baumannii* – either the Pasteur scheme (22) (as used in this study), or the Oxford scheme (23) – to investigate the relatedness of isolates. Where studies have used MLST, the most commonly identified clonal complex is CC^PAS^2 (CC^OX^208). However, a considerable proportion of isolates are often found to belong to less common clonal complexes, or are singletons (16, 19, 27, 63). This is entirely consistent with the results from our study, where 44% of isolates belonged to CC^PAS^2, 24% of isolates belonged to CC^PAS^1, and 19% belonged to ST^PAS^158. The core genome analysis we conducted demonstrated that even within MLST STs there was a lot of diversity. This shows that multiple carbapenem-resistant strains are present within AMUH, suggesting that rather than facing an outbreak, the bacterium is endemic. Whether patients are acquiring these strains once admitted to the hospital, or whether there is widespread circulation of carbapenem-resistant *A. baumannii* in the community is an open question that we hope to address in the future.

While CC^PAS^1 and CC^PAS^2 strains are globally distributed and frequently encountered, strains belonging to ST^PAS^158 have been reported far less frequently and from a more focused geographic area. ST^PAS^158 belongs to CC^PAS^158 (CC^OX^499) (64) and is usually found in isolates from North Africa, the Middle East and Pakistan (Table 2). Most previous studies that have identified CC^PAS^158 isolates have found them to carry the OxaAb variant OxaAb(65). However, in CC^PAS^158 strains the *oxaAb*(65) allele differs from the original allele (accession no. AY750908) by three synonymous substitutions. As the *oxaAb* genes are intrinsic to *A. baumannii* and specific alleles are associated with certain international clones (ICs), the gene can be used as a useful epidemiological marker to identify the IC an isolate belongs to (57, 65, 66). However, under this scheme, OxaAb(65) is associated with IC5. Isolates belonging to IC5 are members of CC^PAS^79 and are found at particularly high frequency in Latin America (15, 57, 67, 68). The allele profiles of the founder sequence types of CC^PAS^158 and CC^PAS^79 (ST^PAS^158 and ST^PAS^79 respectively) are quite different, sharing only 1 of the 7 alleles (*rplB* allele 4), which at the nucleotide level translates to 13 SNPs. It is therefore clear that in this instance, numbering the *oxaAb* alleles based upon their amino acid sequence can mask important epidemiological information and that, as suggested by Karah *et al* (64), these genes should be numbered according to their nucleotide sequences as has been done for the *Acinetobacter ampC* genes (69).

Previous studies of *A. baumannii* in Egypt have found that rates of carbapenem resistance are high, typically >70% (17, 70), and that this is usually associated with isolates carrying the *oxa23* gene with carriage frequencies reaching as high as 100% in carbapenem-resistant isolates (16–18). This was reflected in our study, where *oxa23* was carried by 100% of carbapenem-resistant isolates. Reports of the metallo-β-lactamases NDM-1 and NDM-2 being encoded by isolates from Egypt indicate frequencies of *bla*_NDM-1_ can typically reach up to 30% (18, 71, 72), though reports from specific hospitals can occasionally report higher frequencies (16, 73). This is in line with our study where 6 isolates (11%) carried a *bla*_NDM-1_ gene and only 1 isolate (2%) carried a *bla*_NDM-2_ gene. It is possible that the almost ubiquitous nature of the *oxa23* gene has reduced the selective advantage for subsequent acquisition and retention of *bla*_NDM_ genes, limiting their spread within *A. baumannii*. The *oxa23* gene in *A. baumannii* is typically carried on a transposon mobilised by insertion sequences (ISs), usually IS*Aba1* (10). The IS elements are located immediately upstream of the *oxa23* gene, where they provide a promotor sequence that drives high level expression of *oxa23* (13, 58). The most commonly reported transposons carrying *oxa23* are *Tn2006* which is a composite transposon where *oxa23* and three other genes are bracketed by two IS*Aba1* elements (58), and *Tn2008* which is a one-ended transposon with a single IS*Aba1* element upstream of the *oxa23* gene (74). While the limitations of short-read sequencing in enabling the assembly of transposons is well known, in our study we were nevertheless able to identify a large number of different genetic arrangements surrounding the *oxa23* gene. In line with what is reported in the literature, a structure likely to be *Tn2006* was the most common arrangement in our isolates. However, the large number of different structures we have identified involving *ISAba125, ISAba33* and *ISAba4* in addition to *ISAba1* demonstrate that the carbapenem-resistant *A. baumannii* population in AMUH is not dominated by a single plasmid that is disseminating *oxa23*. Rather, a multitude of different mobile elements are hosting the gene, consistent with the apparent endemic nature of *oxa23* in the bacterial population.

The carriage of antibiotic resistance genes on transposons is common, and is the typical genetic context for OXA-type carbapenemases in *A. baumannii*. However, in isolates belonging to CC^PAS^1 in our study the most commonly identified mobile element carrying *oxa23* was a bacteriophage phiOXA. Reports of the carriage of antibiotic resistance genes in prophages have become more common in recent years (75–77), but it is thought that this is generally a rare occurrence (78). However, recent evidence from studies focusing on *A. baumannii* have suggested that carriage of both virulence and antibiotic resistance genes by prophages is relatively common in this species and may be a major mechanism of horizontal transfer of these genes (79–83). It was recently noted that prophages appeared to be more common in IC5 isolates than in those belonging to IC1 or IC2 (81) and it is an intriguing possibility that prophages may be a major factor in the evolution of different international clones. The carriage of OXA-type carbapenemases in prophages has been observed previously, with *oxa58* identified on a prophage in a *Proteus mirabilis* strain (84) and *oxa23* identified on a prophage in *A. baumannii* strain ANC 4097 (79, 80) and on the phage phiOXA in isolate AB5075-UW (59). However, López-Leal *et al* (85) recently indicated that OXA carbapenemases in prophages may be more widespread, with evidence for potential OXA prophage carriage in approximately 25% of isolates studied. Similarly, we found *oxa23* carried on phiOXA in 15% of our isolates. Moreover, these isolates were not clonally related within CC^PAS^1 but were spread throughout the CC^PAS^1 clade, indicating that phiOXA is widely disseminated amongst CC^PAS^1 isolates in AMUH. Furthermore, we demonstrated that phiOXA can be induced from multiple isolates and that the induced phage particles are carrying the *oxa23* gene. It is therefore clear that in *A. baumannii* bacteriophages can be a major mechanism for the mobilisation of antibiotic resistance genes including those of greatest clinical concern such as the carbapenemases. As more genomic studies using long-read sequencing are conducted that can properly resolve complex mobile element structures, the true magnitude of bacteriophage-mediated antibiotic resistance gene carriage will be revealed.

## Authors and contributors

BAE and AA were involved in the conceptualisation, methodology, investigation, writing (original draft preparation, review and editing) and funding of the study. JM and DT were involved in the methodology, investigation and writing (original draft preparation, review and editing) of the study. EL was involved in the investigation and writing (review and editing) of the study.

