## Supplementary figures and images for "Diversity of carbapenem-resistant *Acinetobacter baumannii* and bacteriophage-mediated spread of the Oxa23 carbapenemase"

### Supplementary Figure 1

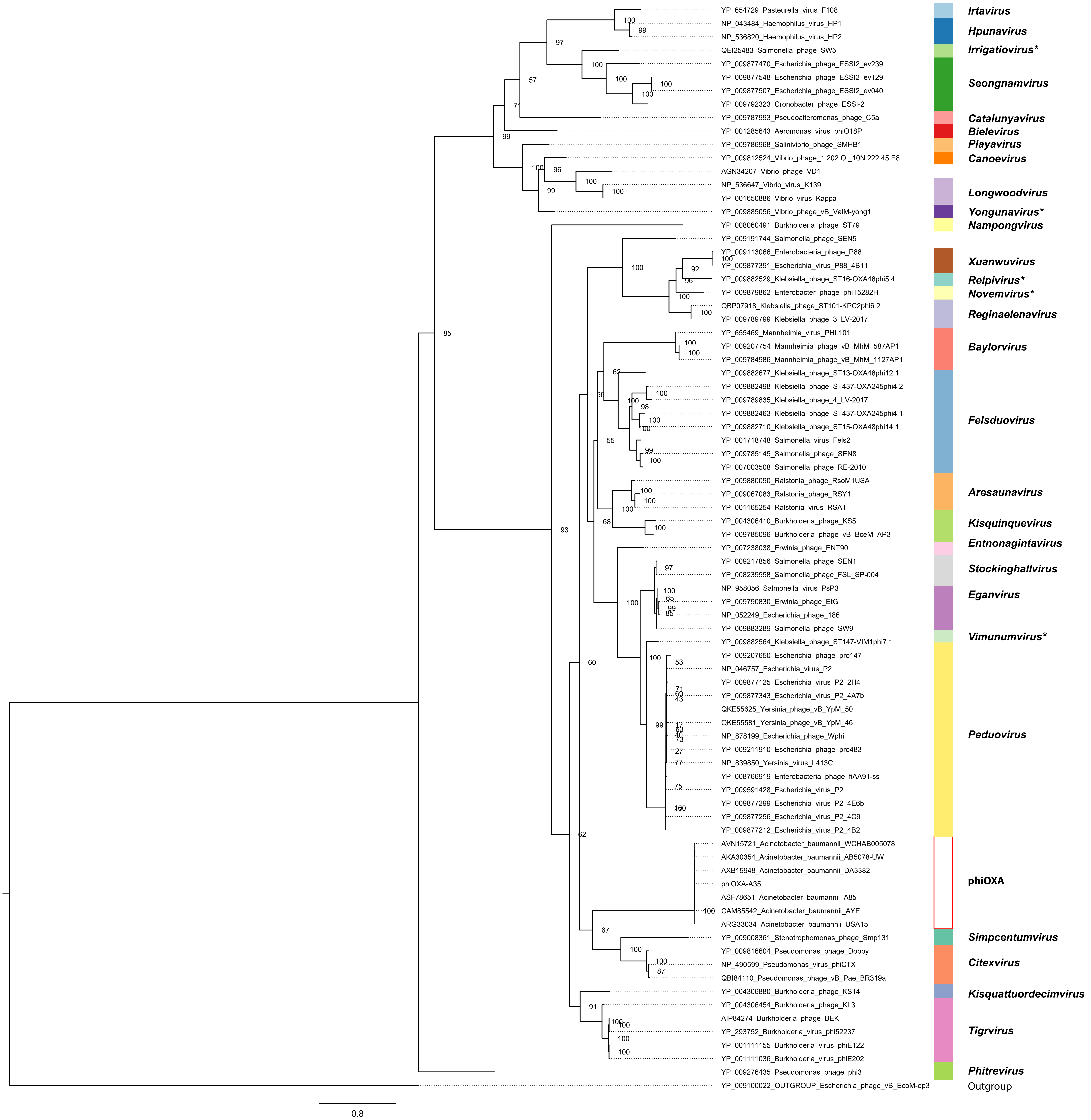

### Supplementary Figure 2

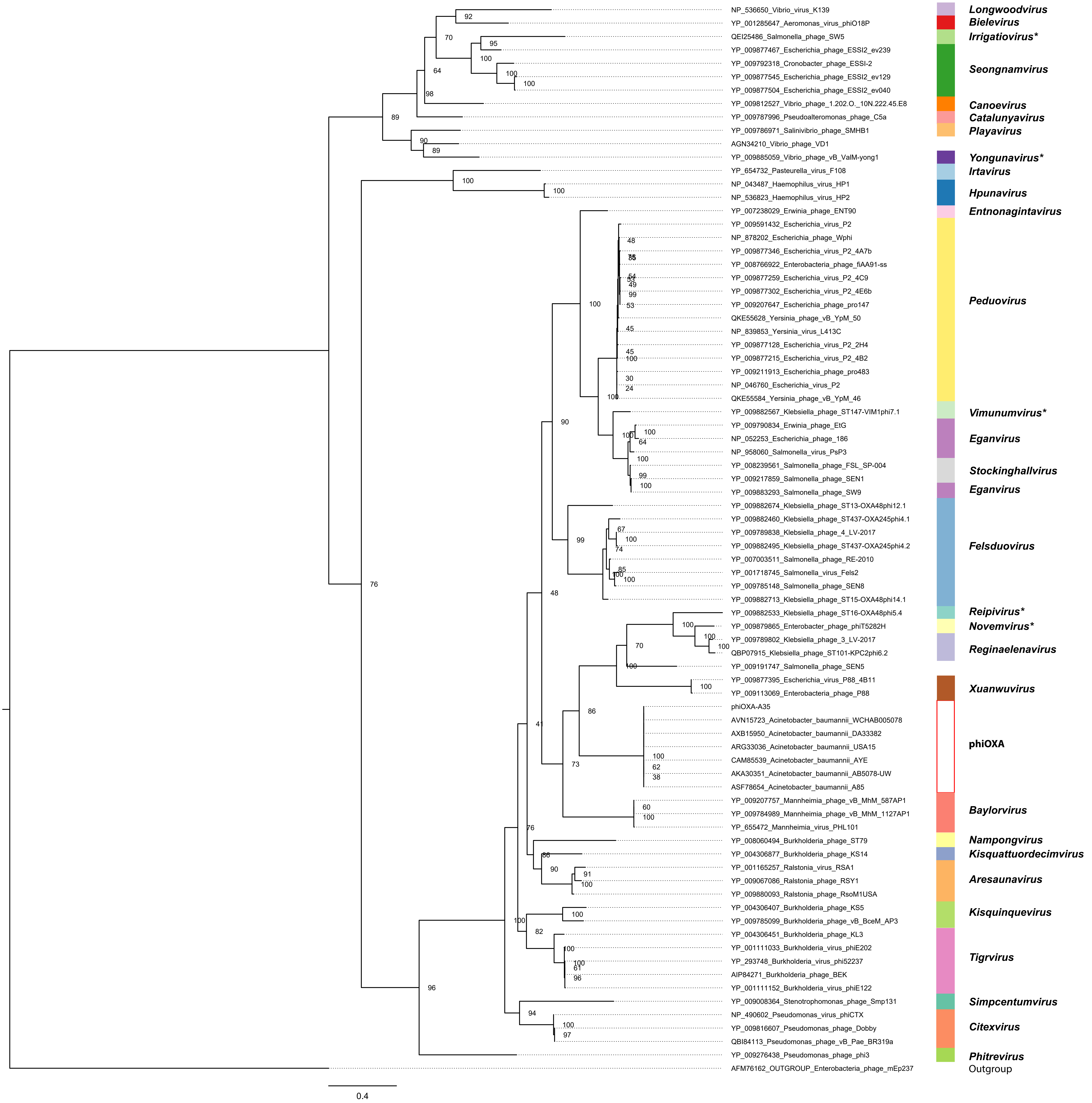
