## Supplementary Figure Legends for "Diversity of carbapenem-resistant *Acinetobacter baumannii* and bacteriophage-mediated spread of the Oxa23 carbapenemase"

Figure S1. Maximum-likelihood phylogenetic tree created from portal vertex protein sequences of members of the subfamily *Peduovirinae*. Sequences were aligned with Clustal Omega and trees constructed using IQTree v1.6.12 with the LG + G4 substitution model IQTree v1.6.12 with ModelFinder, SH-aLRT test and ultrafast bootstrap (1000 replicates). Enterobacteria phage mEp237 (JQ182730) was used as an outgroup to root the tree. Branch length is proportional to the number of substitutions per site (see scale bar). Members of virus genera are denoted by coloured blocks. An asterix (*) adjacent to a genus name indicates proposed genera that have yet to be ratified by the International Committee on the Taxonomy of Viruses [1].

Figure S2. Maximum-likelihood phylogenetic tree created from major capsid protein sequences of members of the subfamily *Peduovirinae*. Sequences were aligned with Clustal Omega and trees constructed using IQTree v1.6.12 with the WAG + G4 substitution model, SH-aLRT test and ultrafast bootstrap (1000 replicates). Enterobacteria phage mEp237 (JQ182730) was used as an outgroup to root the tree. Branch length is proportional to the number of substitutions per site (see scale bar). Members of virus genera are denoted by coloured blocks. An asterix (*) adjacent to a genus name indicates proposed genera that have yet to be ratified by the International Committee on the Taxonomy of Viruses [1].
